# A Novel Method to Visualize Active Small GTPases Unveils Distinct Sites of Sar1 Activation During Collagen Secretion

**DOI:** 10.1101/2024.03.20.586037

**Authors:** Miharu Maeda, Masashi Arakawa, Yukie Komatsu, Kota Saito

## Abstract

Small GTPases are essential in various cellular signaling pathways, and detecting their activation within living cells is crucial for understanding cellular processes. Fluorescence resonance energy transfer is widely used to study the interaction between activated small GTPases and their effectors, but it is limited to those with well-defined effectors, excluding Sar1. Here, we present a novel method, SAIYAN (Small GTPase ActIvitY ANalyzing), for detecting the activation of endogenous small GTPases via fluorescent signals utilizing a split mNeonGreen system. We demonstrated Sar1 activation at the endoplasmic reticulum (ER) exit site and successfully detected its activation state in various cellular conditions. Utilizing the SAIYAN system in collagen-secreting cells, we discovered activated Sar1 localized both at ER exit sites and ER-Golgi intermediate compartment (ERGIC) regions. Additionally, impaired collagen secretion led to the confinement of activated Sar1 at the ER exit sites, underscoring the significance of Sar1 activation through the ERGIC in collagen secretion.

## INTRODUCTION

Small GTPases, cycling between inactive guanosine diphosphate (GDP) and active guanosine triphosphate (GTP) forms, serve as molecular switches in various cellular processes, including cell proliferation, growth, differentiation, motility, and membrane trafficking^1, 2, 3^. The activation of these small GTPases is catalyzed by guanine nucleotide exchange factors (GEFs), which destabilize the binding of GDP to GTPase, thereby facilitating its transition to the free form capable of recruiting cytoplasmic GTP. Upon binding to GTP, the protein interacts with effectors to execute cellular functions. Generally, the intrinsic GTPase activity of small GTPases is relatively slow, and inactivation is facilitated by GTPase-activating proteins (GAPs)^4^.

The conventional approach to assessing GTPase activity *in vitro* relies on the use of radioisotopes ^5, 6^. Methods such as differential tryptophan fluorescence measurements of small G proteins in their GDP- and GTP-bound states can be used as potential improvements to these assays for GTPases containing tryptophan residues (excluding the Ras family)^6, 7^. Another approach involves the use of a fluorescent nucleotide such as mant-GTP^8^.

In addition to the *in vitro* assays, the measurement of GTPase activity within cells is essential, as it directly reflects the broad cellular responses mediated by GTPases with effector proteins. There are currently three primary methods for measuring the activity of small GTPases within cells. Initially, ^32^Pi labeling of cells was used to directly measure the ratio of bound nucleotides to small GTPases^9, 10^. Pulse-labeled cells were extracted, followed by immunoprecipitation of small GTPases, and the bound nucleotides were resolved using thin-layer chromatography to determine the proportion of GDP to GTP. Although highly sensitive, this assay requires radioisotope labeling, is time-consuming, and lacks the ability to monitor changes over time.

Alternatively, the pull-down assay offers a method for measuring activity^11, 12^. In this method, cells are extracted, activated small GTPases are captured by effector proteins, and the number of bound GTPases is quantified using western blotting. Despite being relatively straightforward, this assay still requires cell extraction, and GTPase hydrolysis needs to be taken into account during experiments.

The third assay utilizes fluorescence resonance energy transfer (FRET)-based biosensors^13^ to transfect cells, enabling the timely measurement of GTPase activity within living cells. The Raichu, Ras superfamily, and interacting protein chimeric unit represent the first intramolecular biosensors designed to detect FRET between YFP and CFP upon Ras activation and its interaction with the effector Raf^14^. Although this method has been successfully employed for various GTPases, it is limited to GTPases with well-defined effector molecules. Moreover, the overexpression of FRET sensors in cells restricts the monitoring of endogenous protein activity to specific cellular locations.

Split fluorescent systems offer versatile tools for a wide array of applications, including systematic endogenous tagging for visualizing endogenous protein localization and detecting membrane contact sites^15, 16, 17, 18^. In this study, we used this system to measure the activation of endogenous small GTPases in living cells. Our approach allows detection of Sar1 activation under various physiological locations and conditions. Furthermore, this novel system has potential for application across a broad spectrum of different GTPases, and its effectiveness is discussed in subsequent sections.

## RESULTS

### Validation of SAIYAN technology

Sar1 is a small GTPase with an amphipathic N-terminus, belonging to the Arf-family^19^. Upon activation by the guanine nucleotide exchange factor (GEF) Sec12, the N-terminus of Sar1 gets inserted into the endoplasmic reticulum (ER) membrane to facilitate the recruitment of Sec23/Sec24^20, 21, 22^. Sec23 acts both as an effector by interacting with activated Sar1 and as a GTPase-activating protein (GAP) for Sar1^23^. This dual role poses challenges for the development of conventional FRET-based sensors to monitor Sar1 activity.

To detect Sar1 activation within living cells, we initially established stable cell lines expressing 10 of the 11 bundles of mNeonGreen2 (mNG_1-10_) tethered to the ER membrane in HeLa cells, which were obtained by fusion with the membrane-spanning region of TANGO1S and an HA tag (Fig. 1a). This cell line was named HA-mNG_1-10_. Following doxycycline induction, we verified that the constructs labeled with HA were distributed throughout the ER, exhibiting a characteristic reticular staining pattern that colocalized with PDI (Supplementary Fig. 1a, upper panel). However, this construct did not produce any mNeonGreen2 (mNG) signal (Supplementary Fig. 1a, upper panel). Next, we transiently expressed Sar1A constructs containing a FLAG tag, followed by a glycine linker fused to the 11th bundle of mNG (Sar1A-FLAG-mNG_11_) in HA-mNG_1-10_ cells (Fig. 1a). No mNG signal was detected in the absence of HA-mNG_1-10_ induction (Supplementary Fig. 1b, upper panel). However, upon induction of HA-mNG_1-10_, mNG signals became apparent (Fig. 1b, upper panel; Supplementary Fig. 1a, bottom panel; Supplementary Fig. 1b, bottom panel). These mNG signals exhibited a pattern distinct from those of HA or PDI, appearing as punctate structures in the ER membrane network (Supplementary Fig. 1a, bottom panel). Costaining with Sec16, a well-established ER exit site marker, revealed mNG localization at these sites (Fig. 1b, upper panel)^24^. Notably, FLAG staining, representing the entire Sar1A protein irrespective of its nucleotide status, was distributed in both the cytoplasm and ER exit sites, maintaining its localization regardless of the expression of HA-mNG_1-10_ (Supplementary Fig. 1b). This observation indicated that Sar1A-FLAG-mNG_11_ was not artificially recruited to the ER membrane by mNG_1-10_ to form complete mNG proteins. Therefore, we inferred that the mNG signals corresponded to activated Sar1A and proceeded to further validate this methodology by examining the activation of various Sar1A mutants.

**Fig. 1:**
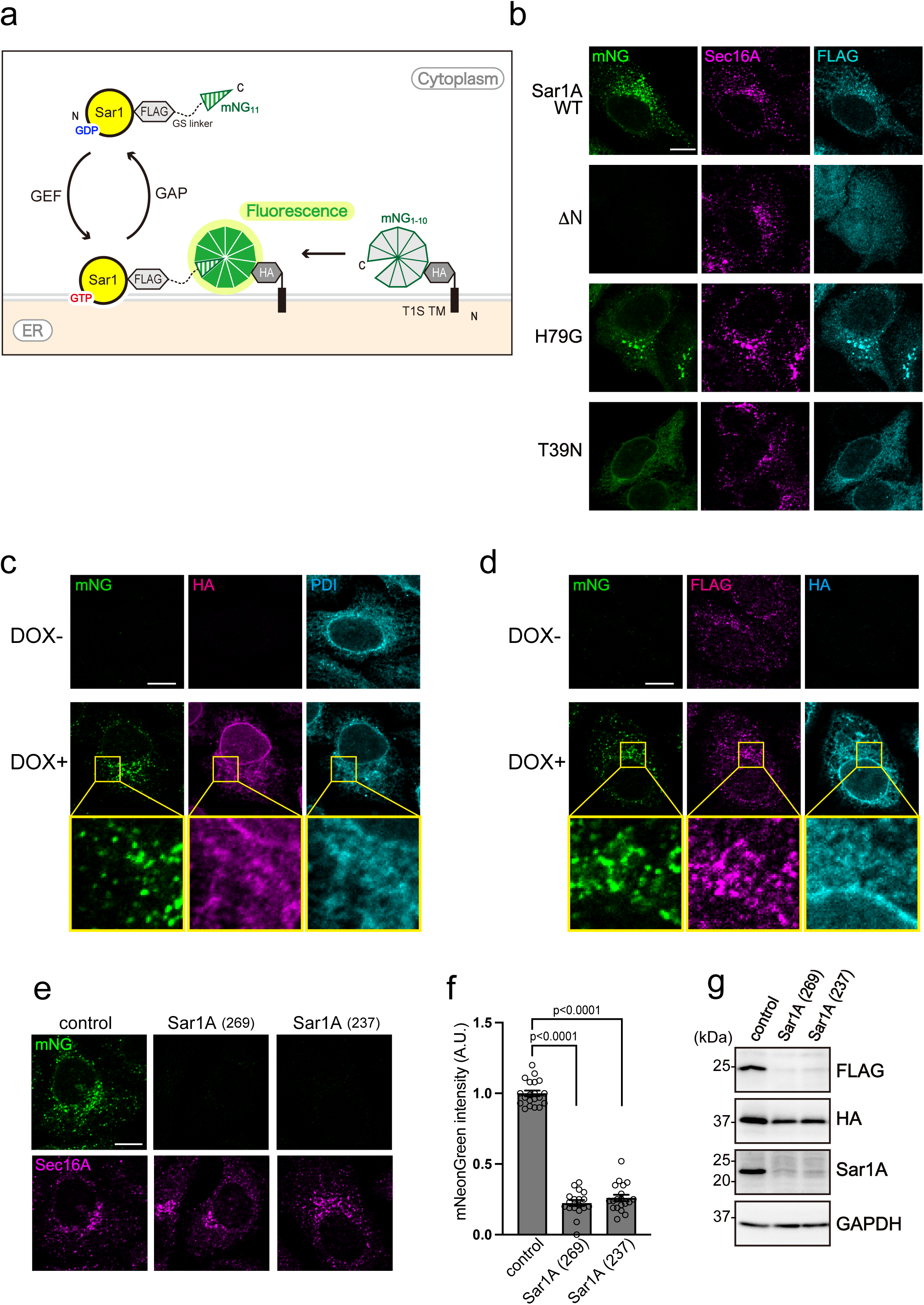
Production of Sar1A/SAIYAN cells. **a** Schematic representation of SAIYAN technology. The membrane-spanning regions of TANGO1S and HA-tag fused to 10 of the 11 bundles of mNG (mNG_1-10_) were expressed in cells. In addition, Sar1A constructs with a FLAG tag and a glycine linker fused to the 11th bundle of mNG (mNG_11_) were also expressed. Upon Sar1A activation, mNG_1-10_ and mNG_11_ combine to form the complete mNG proteins, inducing mNG signals. **b** HA-mNG_1-10_ cells transfected with the indicated Sar1A constructs were fixed and stained with anti-Sec16-C and anti-FLAG antibodies. Scale bars = 10 µm. **c** Sar1A/SAIYAN (HeLa) cells, either treated or non-treated with doxycycline, were fixed and stained with anti-HA and anti-PDI antibodies. Boxed areas in the middle panels are shown at high magnification in the bottom panels. Scale bars = 10 µm. **d** Sar1A/SAIYAN (HeLa) cells, either treated or non-treated with doxycycline, were fixed and stained with anti-HA and anti-FLAG antibodies. Boxed areas in the middle panels are shown at high magnification in the bottom panels. Scale bars = 10 µm. **e** Sar1A/SAIYAN (HeLa) cells transfected with the indicated siRNAs were fixed and stained with an anti-Sec16-C antibody. Scale bars = 10 µm. **f** Quantification of mNG intensity from (**e)**. (arbitrary units [A.U.]). Error bars represent the means ± SEM. **g** Sar1A/SAIYAN (HeLa) cells transfected with the indicated siRNAs were lysed, and subjected to SDS-PAGE, followed by western blotting with anti-FLAG, anti-HA, anti-Sar1A and anti-GAPDH antibodies.

The ΔN mutant, which fails to insert into the ER membrane, did not generate any mNG signals despite expressing Sar1A at a level comparable to that of the wild-type Sar1A, as indicated by FLAG staining (Fig. 1b)^25^. This observation further supports the notion that mNG signals reflect the activation status of Sar1A at the ER membrane. Consistent with this finding, the T39N mutant, deficient in GTP binding, exhibited weak signals throughout the ER, underscoring the necessity of Sar1A activation for robust mNG signaling (Fig. 1b)^26^. In contrast, the Sar1A H79G mutant, a GTPase-deficient variant of Sar1A, generated strong mNG signals that colocalized with Sec16 (Fig. 1b)^27^. The localization of mNG signals in various mutants was consistent with previously reported biochemical properties of Sar1 ^19^. Consequently, we conclude that the system can detect the activation of small GTPases in living cells. We named this technology Small GTPase ActIvitY ANalyzing (SAIYAN). Subsequently, we established cell lines capable of detecting endogenous Sar1 activation. Endogenous Sar1A was tagged with mNG_11_ using CRISPR/Cas9-mediated knock-in tagging of HA-mNG_1-10_ cells (Supplementary Fig. 2). These cell lines were validated by sequencing the Sar1A genomic locus and designated as Sar1A/SAIYAN (HeLa) cells. Consistent with the transient expression, mNG signals were observed only upon doxycycline induction and were exclusively localized at the ER exit sites (Fig. 1c, d). Using siRNAs suppressed Sar1 expression in Sar1A/SAIYAN (HeLa) cells, thereby diminishing mNG fluorescence (Fig. 1e). Quantification of the mNG signals is presented in Fig. 1f. Western blotting of cell lysates confirmed the reduction in bands corresponding to Sar1A and FLAG (Fig. 1g), further validating the production of Sar1A/SAIYAN (HeLa) cells. Moreover, no significant difference in growth of Sar1A/SAIYAN (HeLa) cells was observed compared to the parental HeLa cells, both with and without doxycycline induction, indicating that the constructs were non-toxic to the cells (Supplementary Fig. 3a). Additionally, GM130 staining revealed a typical Golgi structure, further confirming that the constructs had no effect on secretory pathways (Supplementary Fig. 3b).

### Sar1 activation is mediated by cTAGE5/Sec12 at ER exit sites in living cells

The effects of Sec12 depletion were examined. Consistent with our expectations, knockdown of Sec12 using siRNAs resulted in diminished mNG signals (Fig. 2a, b; Supplementary Fig. 4a), verifying the role of Sec12 in mediating Sar1 activation at ER exit sites^20^. Our previous investigations demonstrated that the interaction between Sec12 and cTAGE5 is crucial for the appropriate localization of Sec12 to ER exit sites^28^ and the depletion of cTAGE5 disperses Sec12 throughout the ER^29^. Therefore, the effect of cTAGE5 depletion on Sar1 activity was examined. The depletion of cTAGE5 decreased Sar1 activity at the ER exit sites (Fig. 2c, d), while the expression level of Sec12 remained unchanged (Supplementary Fig. 4b). These findings indicate that the accurate positioning of Sec12 at the ER exit sites is essential for the effective activation of Sar1 at these sites.

**Fig. 2:**
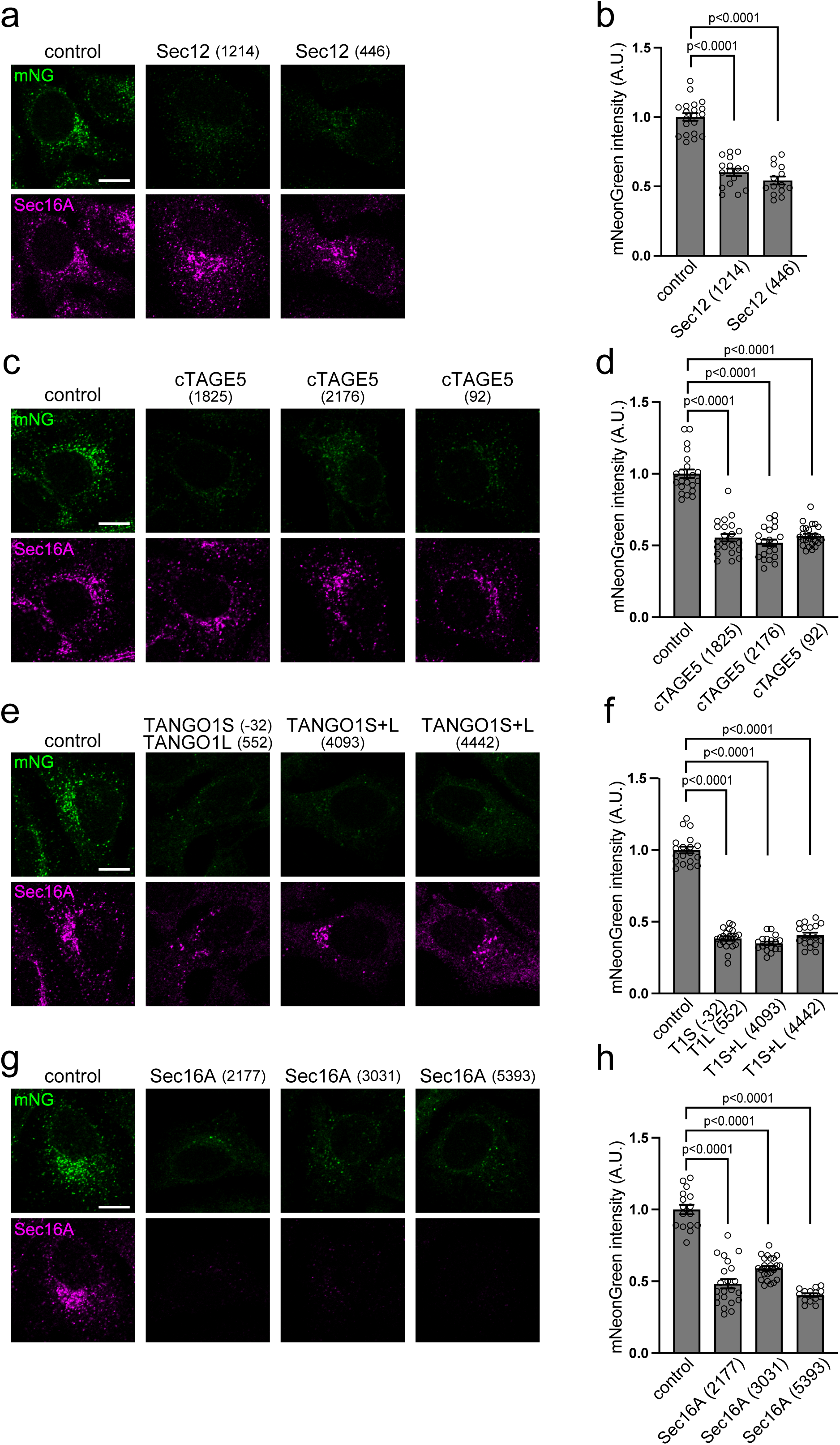
ER exit site organization is required for the efficient activation of Sar1A. **a, c, e, g** Sar1A/SAIYAN (HeLa) cells transfected with the indicated siRNAs were fixed and stained with anti-Sec16-C antibodies. Scale bars = 10 µm. **b, d, f, h** Quantification of mNG signals from (**a), (c), (e), (g),** respectively (arbitrary units [A.U.]). Error bars represent the means ± SEM.

### ER exit site organization is crucial for proper Sar1 activation in living cells

We investigated the effects of perturbing the organization of the ER exit sites on Sar1 activity. Our previous study demonstrated the interaction between TANGO1 and Sec16 and its significance in ER exit-site organization^30, 31^. Consequently, we depleted TANGO1 and Sec16, and examined their effects on Sar1 activity. The knockdown of both TANGO1 and Sec16 resulted in reduced mNG signaling, indicating the essential role of TANGO1 and Sec16 in facilitating Sar1 activation (Fig. 2 e–h; Supplementary Fig. 4c, d).

### Sec23 acts as a stabilizer of activated Sar1 rather than an inactivator in living cells

Our findings suggest that the absence of crucial ER exit-site components results in Sar1 inactivation. To assess whether the SAIYAN system could also detect an increase in Sar1 activity, Sec23A was depleted. Given that Sec23 promotes GTP hydrolysis of Sar1 *in vitro*, its depletion should theoretically enhance Sar1 activity. However, Sar1 activity was severely diminished in Sec23A-depleted cells (Fig. 3 a, b; Supplementary Fig. 5a). Subsequently, we knocked down Sec31A, which is known to enhance Sec23 GAP activity towards Sar1 by approximately ten-fold *in vitro*^7, 32^. As anticipated, Sec31A depletion increased Sar1 activity (Fig. 3c, d; Supplementary Fig. 5b). These results suggest that the SAIYAN system can detect the augmentation of small GTPases. Moreover, we confirmed in living mammalian cells that Sec23 serves as a stabilizer of activated Sar1 rather than an inactivator, as previously suggested by structural and biochemical analyses conducted *in vitro*^32^.

**Fig. 3:**
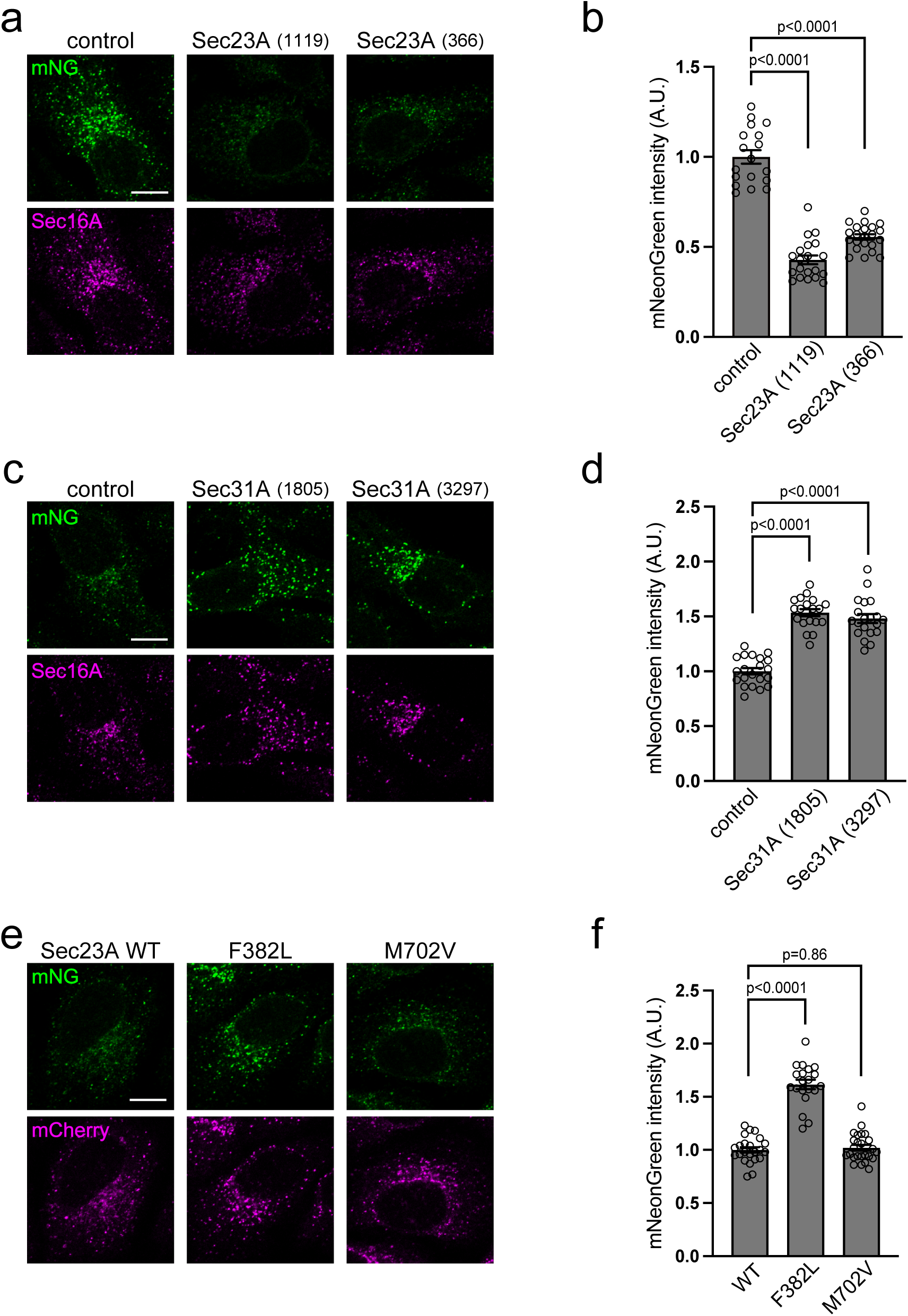
Sec23A and Sec31A depletion exert opposite effects on the activation of Sar1A in living cells. **a, c** Sar1A/SAIYAN (HeLa) cells transfected with the indicated siRNAs were fixed and stained with anti-Sec16-C antibodies. Scale bars = 10 µm. **b, d** Quantification of mNG signals from (**a) and (c),** respectively (arbitrary units [A.U.]). Error bars represent the means ± SEM. **e** Sar1A/SAIYAN (HeLa) cells stably expressing mCherry-tagged Sec23A constructs as indicated were fixed and processed for microscopic analysis. Scale bars = 10 µm. **f** Quantification of mNG signals from **(e)** (A.U.). Error bars represent the means ± SEM.

### CLSD mutants of Sec23A showed activity against Sar1, consistent with *in vitro* studies

Cranio-lenticulo-sutural dysplasia (CLSD; OMIM #607812) is a syndrome characterized by facial dysmorphism, late-closing fontanels, congenital cataracts, and skeletal defects caused by monoallelic or biallelic mutations in the Sec23A gene^33, 34^. Two mutations, F382L and M702V, have been extensively analyzed. While mutation in both alleles causes the F382L mutation to induce CLSD, M702V causes the condition even when only one allele is mutated. Discrepancies in the activities of both mutants towards Sar1A were investigated *in vitro.* Although neither mutation significantly affected Sar1A incubated as Sec23A/Sec24D in the absence of the Sec13/Sec31 complex, the presence of Sec13/Sec31 resulted in a decrease in GAP activity towards Sar1A, particularly in the presence of the F382L mutation^35, 36^. Therefore, using the SAIYAN system, we examined the impact of both mutations on living cells, that contained intrinsic Sec13/Sec31. Stable expression of various mCherry-tagged Sec23A mutants was achieved in SAIYAN cells; however, endogenous Sec23A expression was reduced, as shown in Supplementary Fig. 5c. Sar1 activity was measured in these cells. mNG signals were significantly stronger in cells expressing the F382L mutation (Fig. 3e). This observation was further supported by the quantitative results, which indicate a notable decrease in GAP activity towards Sar1A in cells expressing the F382L mutation (Fig. 3f). Thus, SAIYAN cells allowed for the effective observation of various effects of Sar1 in live cells.

### Activated Sar1 prevails in the ERGIC region of collagen-secreting cells

Our results suggest that the SAIYAN system can detect Sar1 activity in HeLa cells. Next, we investigated Sar1 activity in collagen-secreting cells. Sar1A/SAIYAN cell lines were established from BJ-5ta cells, hTERT-immortalized fibroblasts of human foreskin origin^37, 38^. Genomic sequencing confirmed the production of SAIYAN cells, which were designated as Sar1A/SAIYAN (BJ-5ta) cells. Cells treated with 2,2’-dipyridyl (DPD), a prolyl hydroxylase inhibitor known to impede collagen folding, exhibited augmented procollagen I accumulation within the ER, indicating collagen production and secretion by the cell lines (Supplementary Fig. 6)^39, 40^.

Subsequently, Sar1A/SAIYAN (BJ-5ta) cells were co-stained with Sec16. The cells were visualized using Airyscan microscopy, showing a 1.4-fold improvement in lateral resolution compared to traditional confocal imaging^41^. Sec16 exhibited punctate localization within the cytoplasm, with a tendency to accumulate near the Golgi apparatus, indicating the characteristic localization of ER exit sites, as observed in HeLa cells (Fig. 4a). Upon observing the mNG signal, the punctate localization showed a definite colocalization with Sec16 (Fig. 4a). Notably, in addition to this pattern, the mNG signal exhibited weak extension in a reticular pattern, with less colocalization with Sec16 (Fig. 4a). To further elucidate the nature of these membranous protrusions, we co-stained mNG signals with various antibodies targeting proteins localized at the ER-Golgi interface. This reticular pattern did not coincide with GM130 (Fig. 4m) or PDI (Fig. 4n), which are markers for the Golgi and ER, respectively. However, these reticular membranes containing mNG signals showed significant colocalization with the ERGIC-53 antibodies (Fig. 4b). Conversely, Rab1A, an ERGIC marker, exhibited limited colocalization with activated Sar1 (Fig. 4o). Additionally, Sec23 (Fig. 4c), Sec24B (Fig. 4d), and p125A (Fig. 4f), which are all inner COPII coat constituents, displayed substantial overlap with the reticular membranes of activated Sar1. Additionally, the localization of Sec24D (Fig. 4e) to the reticular signal in the ERGIC region was relatively weaker, and the degree of colocalization with mNG was not as pronounced as that with other inner coat components (Fig. 4p).

**Fig. 4:**
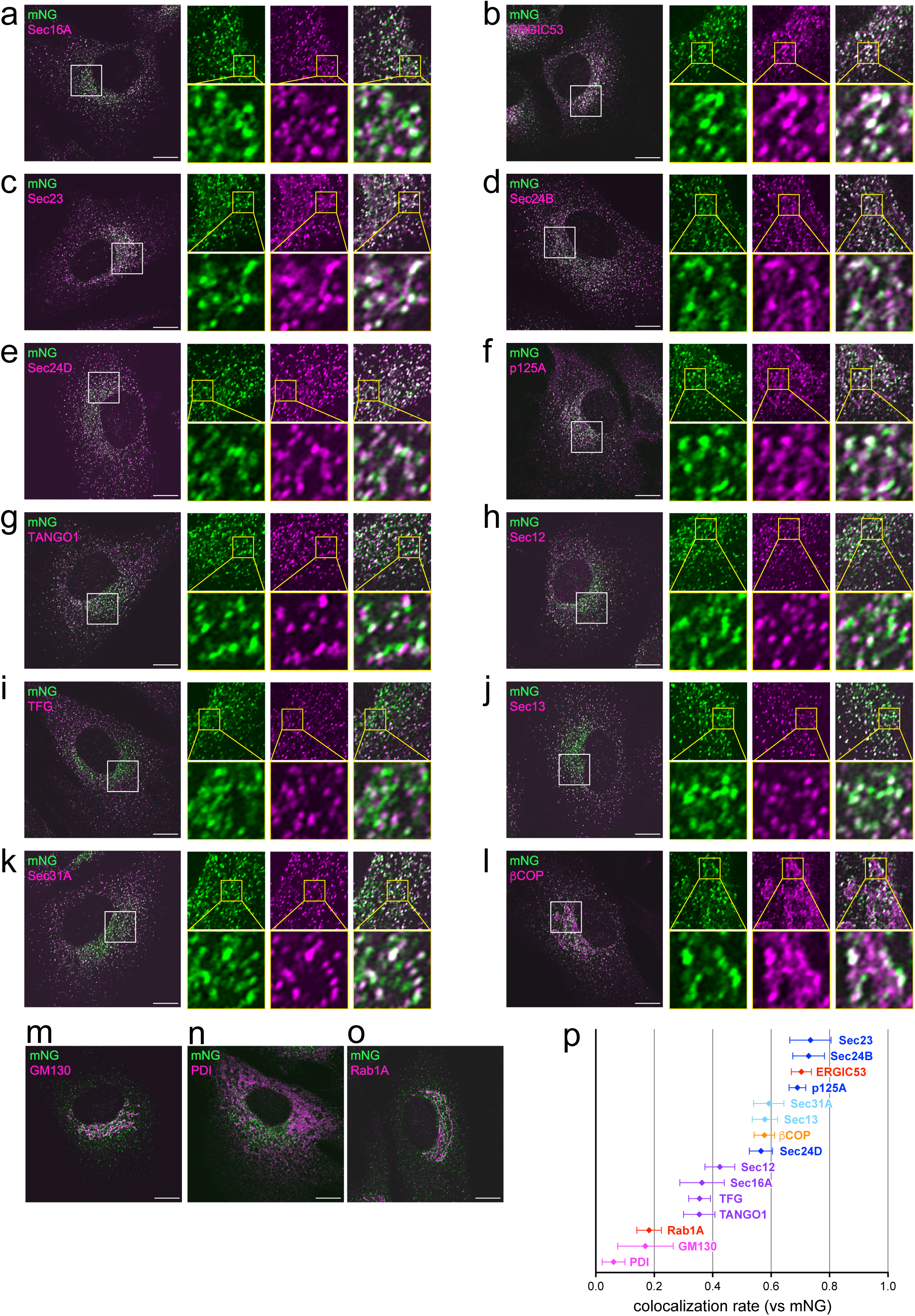
Activated Sar1 prevails in the ERGIC region of Sar1A/SAIYAN (BJ-5ta) cells. Sar1A/SAIYAN (BJ-5ta) cells were fixed and stained with anti-Sec16-C (**a**), anti-ERGIC53 (**b**), anti-Sec23 (**c**), anti-Sec24B (**d**), anti-Sec24D (**e**), anti-p125A (**f**), anti-TANGO1-CT (**g**), anti-Sec12 (**h**), anti-TFG (**i**), anti-Sec13 (**j**), anti-Sec31A (mouse) (**k**), anti-ϕ3-COP (**l**), anti-GM130 (**m**), anti-PDI (**n**), anti-Rab1A (**o**) antibodies. Images were captured using Airyscan2. Scale bars = 10 µm. **a-l** (right; upper) Magnification of the indicated regions is on the left. (right; bottom) Magnification of the indicated regions on the upper. **p** Pearson’s correlation coefficient was used to quantify the degree of colocalization. n=5. cyan; outer COPII coats, blue; inner COPII coats, purple; ER exit site resident proteins, red; ERGIC proteins, orange; COPI protein, magenta; ER and Golgi proteins. Error bars represent the mean 95% CI.

The spatial relationships between these proteins were further elucidated using triple staining, as shown in Fig. 5. As demonstrated by dual staining, Sec16 overlapped with mNG in dotted-like structures, but there is minimal overlap with the reticular patterns of mNG (Fig. 5a). Additionally, as expected, ERGIC53 showed minimal colocalization with Sec16 (Fig. 5a). In contrast, Sec23 not only overlapped with mNG but also colocalized with ERGIC53, demonstrating that Sec23 is localized not only at the ER exit sites but also across the ERGIC region (Fig. 5b). Furthermore, although ERGIC53 and Rab1a partially overlapped, they exhibited distinct regions, suggesting their combined involvement in shaping the ERGIC region (Fig. 5c). mNG signals exhibited considerable overlap with ERGIC53 but minimal overlap with Rab1A, indicating mNG localization to the ERGIC region where ERGIC53 predominates (Fig. 5c).

**Fig. 5:**
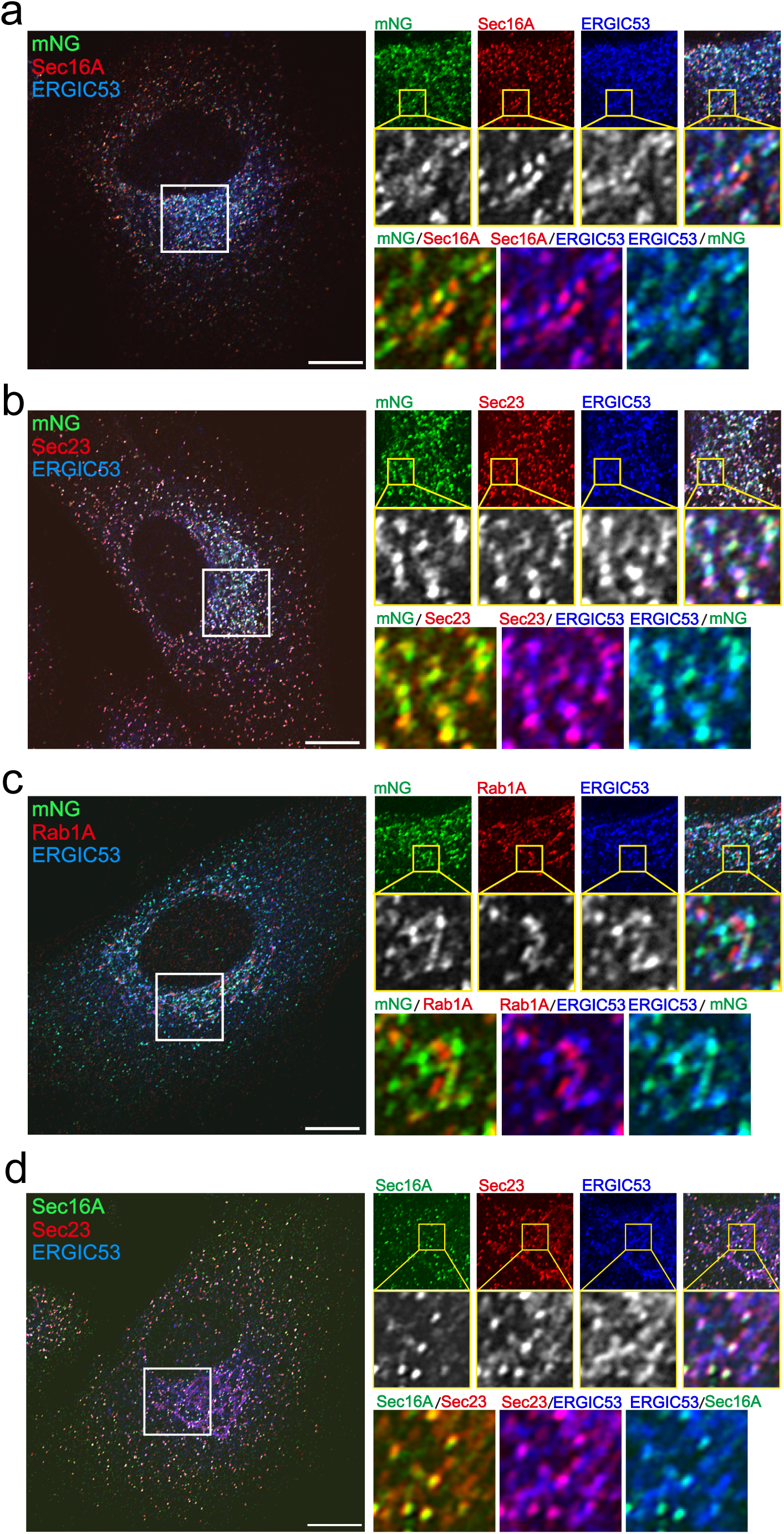
Triple staining of Sar1A/SAIYAN(BJ-5ta) and parental BJ-5ta reveals the organization of the ER-Golgi interface of collagen-secreting cells. **a-c** Sar1A/SAIYAN (BJ-5ta) cells were fixed and stained with anti-Sec16-C and anti-ERGIC53 (**a**), anti-Sec23 and anti-ERGIC53 (**b**), and anti-Rab1A and anti-ERGIC53 (**c**) antibodies. Images were captured using the Airyscan2. Scale bars = 10 µm. **d** BJ-5ta cells were fixed and stained with anti-Sec16-C, anti-Sec23, and anti-ERGIC53 antibodies. Images were captured using the Airyscan2. Scale bars = 10 µm. **a-d** (right; upper) Magnification of the indicated regions is on the left. (right; bottom). Double staining of the magnified region on the upper.

In contrast, ER exit-site proteins, including TANGO1 (Fig. 4g), Sec12 (Fig. 4h), and TFG (Fig. 4i), only exhibited colocalization with dot-like structures similar to those of Sec16. Notable, Sec13 (Fig. 4j) and Sec31A (Fig. 4k), which form the outer COPII coats, exhibited less colocalization with mNG compared to the inner COPII coats, but more than the resident proteins of the ER exit sites. These results indicate that activated Sar1 predominates in the ERGIC regions in conjunction with the inner COPII components within collagen-secreting BJ-5ta cells. Conversely, proteins localized at the ER exit sites remain confined within the contiguous domain of the ER.

Finally, we examined Sec23A localization in parental BJ-5ta cells. As shown in Fig. 5d, Sec23A extension was observed in the ERGIC region, confirming that the reticular signals observed in SAIYAN cells were not a result of introducing the SAIYAN system artificially. This observation further supports the extension of activated Sar1 to ERGIC regions in collagen-secreting BJ-5ta cells.

### The regions of Sar1 activation depend on the cargo secreted by the cell

The aforementioned observation was consistent with previous reports by Weigel et al. and Shomron et al. regarding ER-Golgi protein transport mediated by an interconnected tubular network rather than spherical vesicles, although their reports did not distinguish between resident proteins at ER exit sites and COPII coats ^42, 43^. Furthermore, an intriguing observation in their study was the involvement of COPI proteins in anterograde transport. We investigated the spatial relationship between the COPI proteins and Sar1 activation in SAIYAN cells. Consistent with these findings, COPI proteins were localized in close proximity to the activated Sar1 (Fig. 4l). Fig. 4p illustrates the colocalization rates of activated Sar1 with various factors present at the ER-Golgi interface in Sar1A/SAIYAN (BJ-5ta) cells. Inner COPII components, akin to ERGIC53, exhibit a high degree of colocalization with activated Sar1, whereas outer COPII components and COPI factors subsequently follow. The colocalization rate with resident proteins at ER exit sites, such as Sec12, responsible for activating Sar1, was significantly lower compared to these. A comparison of these results with the colocalization rates of mNG signals with other components present at the ER-Golgi interface in Sar1A/SAIYAN (HeLa) cells would be intriguing. In HeLa cells, the colocalization rate of activated Sar1 was moderately high with ER exit-site resident proteins, similar to inner and outer COPII, whereas ERGIC and COPI exhibited comparatively lower colocalization (Supplementary Fig. 7). These results suggest that the structure of the ER-Golgi interface varies significantly depending on the nature of the secreting cells.

Additionally, to validate this, we treated Sar1A/SAIYAN (BJ-5ta) cells with DPD and examined the change in the activation region of Sar1 upon the inhibition of collagen secretion. Our results revealed that in cells treated with DPD, the previously observed weak reticular signal, which was typical characteristic of BJ-5ta cells, disappeared, and the mNG signal localized to the punctate structures, similar to that observed in HeLa cells (Fig. 6a). In fact, the measurement of colocalization with Sec16 showed a significant increase in DPD-treated cells compared to untreated cells (Fig. 6b). These findings suggest that the ER-Golgi interface adopts different structures depending on the cargo being secreted, thereby maintaining a shape customized for the specific cargo. Furthermore, it became evident that activated Sar1 in collagen-secreting cells was localized along with the inner COPII components from the ER exit site to the ERGIC region.

**Fig. 6:**
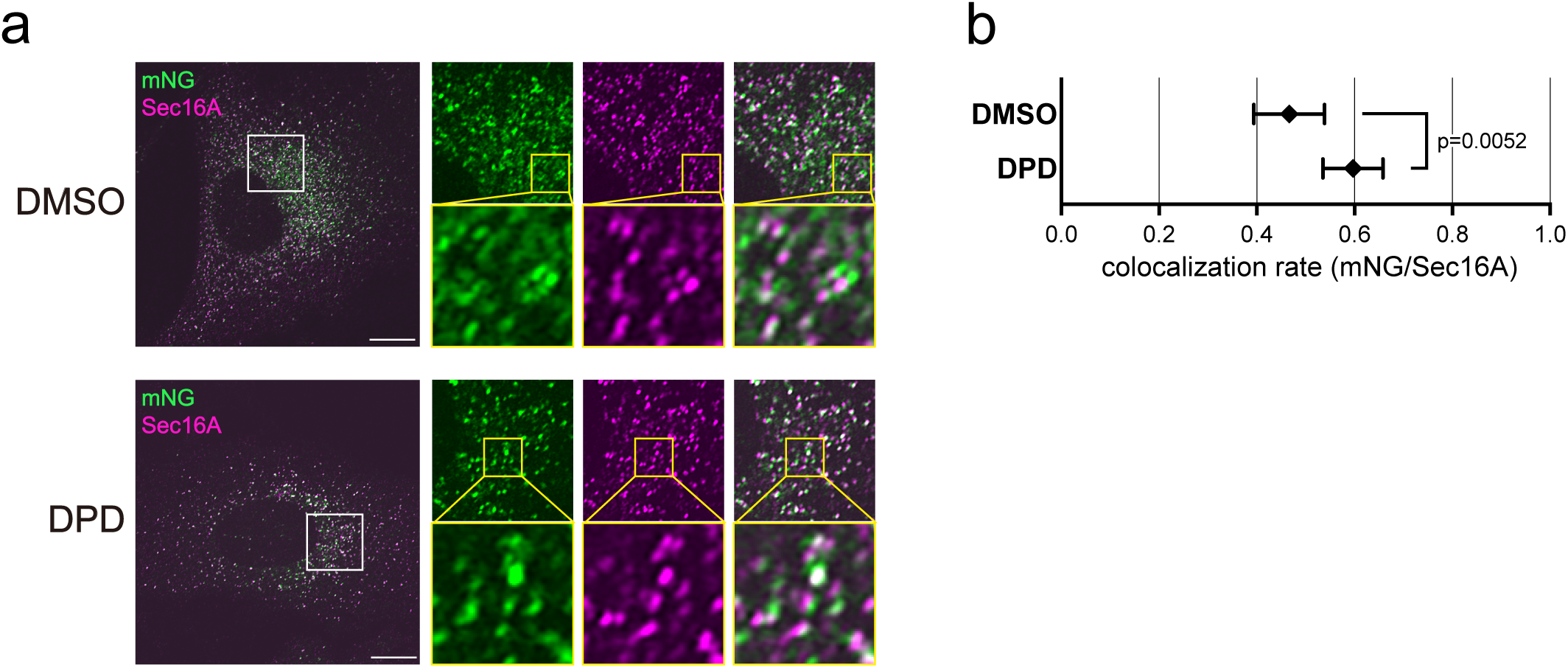
Reticular pattern of activated Sar1 signals diminished with DPD treatment in Sar1A/SAIYAN (BJ-5ta) cells. **a** Sar1A/SAIYAN (BJ-5ta) cells were treated with DMSO or 0.5 mM DPD and incubated for 16 h. Cells were fixed and stained with anti-Sec16-C antibodies. Images were captured using the Airyscan2. Scale bars = 10 µm. **b** Pearson’s correlation coefficient was quantified to assess the degree of colocalization. n=5. Error bars represent the mean 95% CI.

## DISCUSSION

In this study, we successfully visualized the activation of Sar1 in living cells for the first time using Split Fluorescent Protein technology, designated as the SAIYAN system. Previously, the measurement of Sar1 activation relied solely on *in vitro* biochemical assays. However, SAIYAN technology accurately identified the activity of Sar1 modulators in living mammalian cells. Specifically, we directly measured the requirement of Sec12 as a GEF and cTAGE5 as a recruiter for Sec12 in order to activate Sar1 in living cells. Furthermore, we elucidated the roles of essential factors such as TANGO1 and Sec16, which are components of ER exit sites, in facilitating appropriate Sar1 activation. Additionally, we provided evidence in live mammalian cells that while *in vitro* Sec23 exhibits GAP activity towards Sar1, it contributes to the stabilization of activated Sar1 through pre-budding complex formation, consistent with previous analyses of yeast and mammalian proteins^7, 32^. Furthermore, we confirmed that Sec31 possesses GAP-enhancing activity towards Sec23, which results in Sar1 inactivation. We also obtained conclusions consistent with previous *in vitro* results regarding the activity of well-analyzed CLSD-associated mutants of Sec23A, F382L, and M702V on Sar1A. The M702V mutant has been reported to have more substantial effects on Sar1B than on Sar1A^36^. Further development and analysis with SAIYAN cells for Sar1B are anticipated to enhance our understanding of CLSD within cells. We devised a novel system employing SAIYAN technology, allowing real-time monitoring of the activation of small GTPases in live cells that lacked firmly established effectors—a capability unattainable with previous FRET probes.

We have previously demonstrated that Sec12 accumulates at the ER exit site in a TANGO1- and cTAGE5-dependent manner and is necessary for efficient Sar1 activation^28, 30^. cTAGE5 knockdown inhibits Sec12 accumulation and suppresses collagen secretion. Furthermore, we demonstrated that the accumulation of Sec12 at ER exit sites mediated by cTAGE5 can be complemented by the overexpression of wild-type Sar1, indicating the importance of Sar1 activation at ER exit sites for collagen secretion^29^. However, contrary to our expectations, activated Sar1 was not confined solely to the ER exit site but rather extended beyond the ER exit site and extended into the ERGIC region in collagen-secreting cells, a phenomenon not observed in HeLa cells secreting minimal collagen. Notably, upon the addition of DPD, a collagen folding inhibitor, we found that activated Sar1 was specifically localized to the ER exit site, even in BJ-5ta cells. These results strongly suggest that the specific localization of activated Sar1 to the ERGIC region is crucial for collagen secretion. Moreover, the site of Sar1 activation and its function may differ during the process of collagen secretion.

Based on the insights gained from the activation of Sar1 and collagen transport from the ER, reports have indicated that the addition of Sar1 H79G to semi-intact cells results in tubular structures that emanate from the ER^44^. Similar tubular formation has been observed when Sar1 H79G is introduced to artificial liposomes or incubated with non-hydrolyzable GTP analogs such as GTPγS and GMP-PNP^7, 45, 46, 47^. These findings are consistent with the hypothesis that upon Sar1 activation, the amphipathic N-terminal region of Sar1 becomes embedded in the membrane, potentially inducing membrane curvature^22, 48^. Additionally, using cryo-electron microscopy, Zanetti et al. demonstrated that non-hydrolysable Sar1 and COPII components incubated with giant unilamellar vesicles induced the formation of tubes covered by Sec23/Sec24 and Sec13/Sec31^49^. Remarkably, their model predicted that the tubular structures covered with Sec23/Sec24 would recruit fewer Sec13/Sec31 compared to spherical vesicles. Additionally, an enlarged COPII cage suggested a decrease in Sar1-GTP hydrolytic activity compared with conventional COPII coats. We found that upon the secretion of collagen, activated Sar1 extends from the ER exit site to the ERGIC region, where Sec23/Sec24 and p125 colocalize. This observation is consistent with the tubular structures identified by Zanetti et al. upon the addition of GTP-locked Sar1 *in vitro*. Additionally, the co-localization of Sec13/Sec31 with activated Sar1 was lower than that of inner coat factors, but higher than that of ER exit site-resident proteins, thus supporting their proposed model. Mutations in Sec24D (Fig. 4e) cause Osteogenesis Imperfecta (OI), suggesting that Sec24D is involved in collagen secretion^50^. Therefore, the observation that Sec24D is less colocalized with activated Sar1 in ERGIC protrusions in collagen-secreting cells was unexpected. However, evidence suggests that collagen accumulation is not significant in Sec24D-deficient patient skin fibroblasts and that all Sec24 isoforms are involved in collagen transport, indicating the need for further analysis ^50, 51^.

Furthermore, Malhotra et al. proposed the possibility of ERGIC membrane recruitment to TANGO1 located at ER exit sites, suggesting the formation of large carriers necessary for collagen secretion^52^. Subsequently, they proposed a model in which elongated tunnel-like structures facilitate the secretion of collagen from the ER to the Golgi apparatus^53, 54^. Recent observations using live cells and electron microscopy have revealed reticular structures at the ER-Golgi interface, indicating that protein transport may occur through these reticular/tubular structures instead of vesicular transport alone. Nevertheless, these studies primarily focused on the localization of inner and outer COPII proteins as indicative of proteins at the ER-Golgi interface, which may not have clearly distinguished between the resident proteins at the ER exit sites, COPII proteins, and proteins localized at the ERGIC structures. Moreover, most of these analyses were confined to cells such as HeLa, COS7, and CHO, which are known to secrete limited amounts of collagen^42, 43^. Our observations suggest that, at least in collagen-secreting cells, resident proteins at ER exit sites and COPII proteins likely behave differently depending on the cell type. Thus, our current analysis adds some modifications to their observations. A schematic representation of the ER-Golgi interface is shown in Fig. 7. Factors such as TANGO1, Sec16, Sec12, and TFG were localized specifically at the punctate structures commonly referred to as ER exit sites in collagen-secreting cells, with no evident reticular structures directed towards ERGIC (Fig. 7b). Conversely, inner coat proteins composed of Sec23/Sec24 and p125A protruded towards the ERGIC direction, along with activated Sar1, in addition to their localization at conventional ER exit sites (Fig. 7b). Activated Sar1 was restricted to the ERGIC53-positive region of the ERGIC and was absent from the Rab1 side of these cells (Fig. 7b). This observation correlates with a report indicating that Surf4 is present in tubular ERGIC, which is Rab1-positive and ERGIC53-negative^55^. Furthermore, recent discoveries highlighting the interaction of p125 with VPS13B and its role in collagen secretion via tubular ERGIC formation underscore its significance^56^. Moreover, our discovery that activated Sar1 in collagen-secreting cells demonstrates stronger colocalization with β-COP compared to ER exit site proteins holds considerable significance (Fig. 4p). This observation is consistent with prior evidence implicating COPI in collagen secretion^57^. In contrast, in HeLa cells, both resident and COPII proteins were found to colocalize well with activated Sar1 at the ER exit sites, while ERGIC and COPI exhibited less colocalization with activated Sar1 (Fig. 7a).

**Fig. 7:**
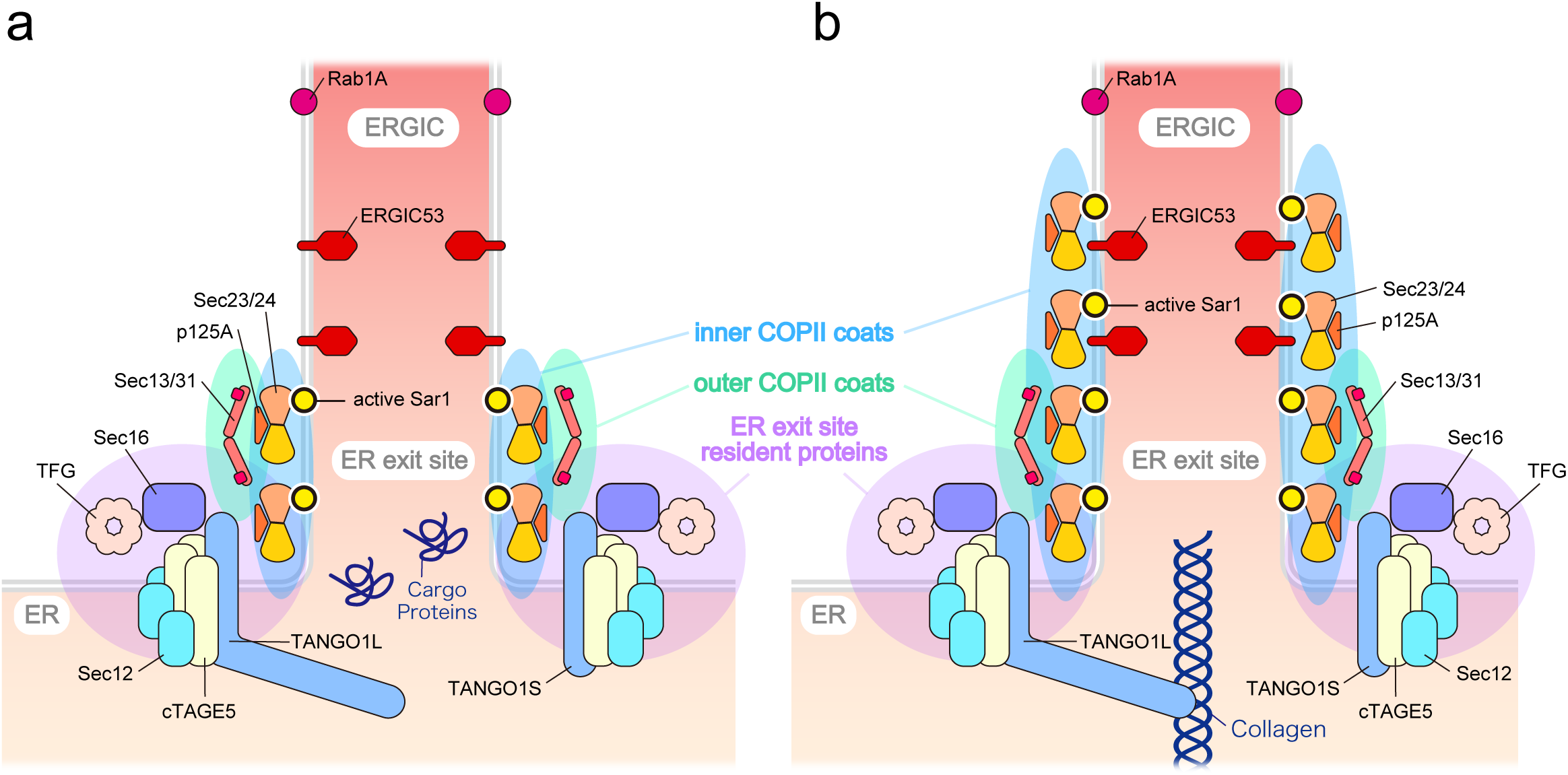
Schematic representation of ER-Golgi interfaces observed with SAIYAN system. **a** In HeLa cells, ER exit site resident proteins exhibit strong colocalization with inner and outer COPII coat proteins at the ER exit sites. In contrast, ERGIC is located away from the ER exit sites. **b** In collagen-secreting BJ-5ta cells, although ER exit site resident proteins are confined to the ER exit sites, inner COPII coats extend to the ERGIC regions labeled with ERGIC-53 but not Rab1. Outer coat proteins partially colocalize with ER exit sites.

Our findings are consistent with both the vesicle-mediated and tunnel-mediated transport models. It remains a challenge to determine whether all secretory proteins are exclusively transported via reticular structures or if both vesicular transport and reticular structures coexist^58^. In this context, whether the differences in the structure of the ER-Golgi interface can be attributed to the mode of secretion is a highly intriguing issue. The mechanism by which activated Sar1 is stabilized and facilitates protrusions into the ERGIC in collagen-secreting cells compared to minimal collagen-secreting cells remains a subject for future investigation. Additionally, variations in the structure of the ER-Golgi interface across different cell types suggest the flexibility of the ER-Golgi interface to accommodate cargo types specific to each cell type. Indeed, in our experiments, treatment of collagen-secreting cells with DPD resulted in their ER-Golgi interfaces resembling those in HeLa cells. Recent studies indicating normal secretion in cells lacking Sar1 and the role of mechanical strain in regulating secretion via Sar1 through Rac1 activity, emphasize the importance of elucidating Sar1’s role in secretion^59, 60^. Thus, understanding the role of Sar1 in vesicular secretion is critical for future research.

One limitation of the current approach is the inherent tendency of split fluorescent proteins to self-associate to some extent^18^. While split fluorescent proteins have been successfully utilized as factors forming membrane contact sites, improper levels of expressed factors can lead to the formation of artificial contact sites ^61^. During the process of initial optimization of the system, when we lengthened the glycine linker attached to Sar1, we observed mNG signals by SAIYAN, even in ι1N Sar1 mutants that were not recruited to the membranes (data not shown). Therefore, the proposed system was carefully validated. The ability to accurately measure both Sar1 inactivation and activation through knockdown of each factor served as the foremost validation of the functionality of the assay system. Therefore, careful validation, particularly to ensure the absence of fluorescent signals in the absence of membrane recruitment, is crucial when employing SAIYAN in other small GTPases for further analysis. Moreover, in future iterations of SAIYAN, the use of more transiently interacting fluorescent probes will be beneficial to measure the kinetics of GTPase turnover instead of monitoring long-term changes using siRNA. Nevertheless, in this study, we successfully detected Sar1 activation in cells for the first time using split fluorescent proteins and gained intriguing insights into the differential activation states of Sar1 across different cell types. By further refining the SAIYAN technology, we anticipate gaining broader insights into the intracellular activation of small GTPases.

## METHODS

### Antibodies

Polyclonal antibody against Sec13 were raised in rabbits by immunization with His_6_-Sec13 and affinity-purified by columns conjugated with GST-Sec13. Polyclonal antibody against Sar1A were raised in rabbits by immunization with Sar1A and affinity-purified by columns conjugated with Sar1A. Polyclonal antibody against TFG were raised in rabbits by immunization with TFG (308-396aa.) and affinity-purified by columns conjugated with corresponding region of TFG. Anti-Sec12 (rat), anti-Sec16-N (rabbit), anti-Sec16-C (rabbit), anti-Sec23A (rat), anti-Sec31A(rabbit), anti-cTAGE5-CC1 (rabbit), anti-TANGO1-CC1 (rabbit) and anti-TANGO1-CT (rabbit) antibodies were produced as described previously ^28, 30, 62, 63, 64^. Sec24D antibody was gifted from Schekman Lab. Other antibodies were as follows: GAPDH (mouse; Santacruz), FLAG (mouse; Sigma-Aldrich or rat; Agilent), HA (rat; Roche), Sec31 (mouse; BD), Sec24B (rabbit; Cell Signaling), p125A (rabbit;proteintech), β-COP (mouse;SIGMA), ERGIC53 (mouse; Santacruz), Rab1a (rabbit;Cell Signaling), GM130 (mouse; BD biosciences), PDI (mouse;abcam).

### Constructs

For transient expression of human Sar1A WT or mutant constructs, human Sar1A cDNA fused with FLAG peptide, GS linker (GGGS), and 11^th^ β strand of mNeonGreen2 to C terminal was cloned into pCMV5 vectors that were gifts from D. Russell (University of Texas Southwestern Medical Center, Dallas, TX) ^65^. Λ1N mutant lacks 25aa of N terminal, which is corresponding to 23aa deletion of yeast constructs ^25^.

### Cell culture and transfection

HeLa cells were cultured in DMEM supplemented with 10% FBS. BJ-5ta cells were cultured in high Glucose DMEM supplemented with Medium 199, 10% FBS, and ascorbic acid phosphate. For transfecting siRNA, Lipofectamine RNAi max (Thermo Fisher Scientific) was used. For single siRNA transfection, reverse transfection protocol was used. In the case of double siRNA transfection, after 24h of the first transfection by reverse transfection, forward transfection was conducted according to the manufacturer’s protocol. For plasmid transfection, Fugene 6 (Promega) or Fugene 4K (Promega) were used. Doxycycline-inducible stable cell lines expressing TM-mNG_1-10_ were made with the lentivirus system described previously^66^.TM-mNG_1-10_ consists of 1–92 aa of TANGO1S (transmembrane region of TANGO1S), HA peptide, and 1–213aa (1∼10th β strands) of mNeonGreen2. Proteins were induced by incubation with 1 µg/ml doxycycline for 24h. Stable cell lines expressing mCherry-Sec23A WT, F382L, and M702V were made with lentivirus by replacing Cas9 in LentiCRISPRv2 with Sec23A cDNAs^67^. 2,2’-dipyridyl was purchased from TCI.

### CRSPR-mediated knock-in

sgRNA was prepared essentially as previously described ^65, 68, 69^. The DNA template for sgRNA was generated by PrimeSTAR GXL DNA Polymerase (TAKARA) by overlapping PCR using a set of three primers: BS7R:5’- AAA AAA AGC ACC GAC TCG GTG C-3’ and ML611R:5’- AAA AAA AGC ACC GAC TCG GTG CCA CTT TTT CAA GTT GAT AAC GGA CTA GCC TTA TTT AAA CTT GCT ATG CTG TTT CCA GCA TAG CTC TTA AAC-3’ and Sar1a_sgRNA_F:5’- CCT CTA ATA CGA CTC ACT ATA GGC AAT ATA CTG GGA GAG CCA GGT TTA AGA GCT ATG CTG GAA-3’. In vitro transcription and sgRNA purification were performed by CUGA7 gRNA Synthesis Kit (NIPPON Gene). To construct the donor plasmid for homology-directed repair, the Sar1 genome template was first amplified from HeLa genomic DNA with following primers (CCGCTCTAGAACTAGTACCCAAATGAGCTCTGGC, CGGTATCGATAAGCTTGCATCAGTATTAAATACACATG) and cloned into pBSIISK(-) by In-Fusion HD cloning Kit (TAKARA). Next, BamHI sequence was inserted into the middle of genomic Sar1 by using following primers (TCCTTGGACGGTGAAAATAAAAGAGTTTTACTTC, TCCGTCAATATACTGGGAGAGCCAGCGG). Then, FLAG peptide, GS linker (GGGS) and 214–229 aa of mNeonGreen2 corresponding to the 11^th^ bundle of mNeonGreen2 were inserted into BamHI locus, to express Sar1A as a fusion with these sequences^65^. The construct was then used as a donor template for ssDNA production by Guide-it Long ssDNA production system v2 (TAKARA). Following primers (GCATCAGTATTAAATACACATG, ACCCAAATGAGCTCTGGCCTCCATATC) with or without 5’ phosphorylation was used for amplifying dsDNA for Strandase reaction to make ssDNA. For CRISPR/Cas9-mediated cell line generation of SAIYAN cells, 1×10^5^ cells of HeLa cells or BJ-5ta cells expressing inducible TM-mNG2_1-10_ were transfected with 7.5 pmol of Cas9 (TAKARA), 7.5 pmol of sgRNA, and 0.64 pmol of ssDNA by Nucleofector II (Lonza) or Neon Transfection System (Thermo Fisher Scientific). Cas9 and sgRNA were incubated at room temperature for 20 min prior to transfection. Electroporated cells were cultured in 24-well plates for 5-7 days and transferred to 6-well plates prior to selection by fluorescence-activated single cell sorting using FACS Melody (BD Biosciences).

### Cell viability assay

Cell viability of cultured cells was quantified using the Cell Counting Kit 8 (DOJINDO Labolatories).

### Western blotting

The experiments were essentially performed as described previously ^28^. Cells extracted with extraction buffer consisting of 20 mM Tris-HCl (pH 7.4), 100 mM NaCl, 1 mM EDTA, 1% Triton X-100, and protease inhibitors were centrifuged at 20,000 g for 15 min at 4°C. The blots were analyzed using ImageQuant LAS4000 or ImageQuant 800 (GE Healthcare).

### siRNA oligos

siRNA oligos used in this study is shown in Supplementary Table 1. The number in the parentheses represents the starting base pair of the target sequence. For control siRNA, Stealth RNAi siRNA Negative Control Med GC Duplex #2 (Thermofisher Scientific) was used.

### Immunofluorescence microscopy

Immunofluorescence microscopy analysis was performed as described previously^28^. Cells grown on coverslips were washed with PBS, fixed with methanol (6 min at −20°C), and then washed with PBS and blocked in blocking solution (5% BSA in PBS with 0.1% Triton X-100 for 15 min). After blocking, cells were stained with primary antibody (1 h at room temperature) followed by incubation with Alexa Fluor-conjugated secondary antibodies (Alexa Fluor 488, Alexa Fluor 568, and/or Alexa Fluor 647 for 1 h at room temperature). Images were acquired with confocal laser scanning microscopy (Plan Apochromat 63×/1.40 NA oil immersion objective lens; LSM 900; Carl Zeiss). The acquired images were processed with Zen Blue software (Carl Zeiss). All imaging was performed at room temperature. mNeonGreen intensity scanning was performed by Fiji-ImageJ^70^. Pearson’s colocalization rate was analyzed by Zen Blue software (Carl Zeiss). Data were analyzed by an un-paired two sample Student’s t-test for two-group comparisons, or a one-way analysis of variance (ANOVA) followed by Dunnett’s test for multiple comparisons using GraphPad Prism 8 software (GraphPad). All graphs were created using GraphPad Prism 8 software (GraphPad).

## Acknowledgements

This work was supported by Grants-in-Aid for Scientific Research (20K15740, 22H02760 to M.M. 23H05254 to Y.K. and 19K22612, 20H03203, 20H04897, 21K19470, 23H02430 to K.S.) from the Ministry of Education, Culture, Sports, Science and Technology of Japan, by Akita University Support for Fostering Research Project to M.M., Y.K., and K.S., by The Naito Foundation to K.S. and M.M., by Takeda Science Foundation to K.S., by Toray Science Foundation (19-6005) to K.S., by The Sumitomo Foundation to K.S., by Suzuken Memorial Foundation to K.S., by The Yasuda Medical Foundation to K.S., by Foundation for Promotion of Cancer Research to K.S., by The Asahi Glass Foundation to K.S., by Princess Takamatsu Cancer Research Foundation to K.S., by Japan Foundation for Applied Enzymology to M.M., by Kato Memorial Bioscience Foundation to M.M., by Astellas Foundation for Research on Metabolic Disorders to M.M., by Inamori Foundation to M.M., by The Kao Foundation for Arts and Sciences to M.M. and by Koyanagi Foundation to M.M.

## Author Contributions

M.M. designed and performed research, analyzed data, and wrote the manuscript; M.A. and Y.K. performed research; and K.S. designed and performed research, analyzed data, and wrote the manuscript.

## Competeing Interest

The authors declare no competing interests.

