## Supplemental Figures and Table for "A Novel Method to Visualize Active Small GTPases Unveils Distinct Sites of Sar1 Activation During Collagen Secretion"

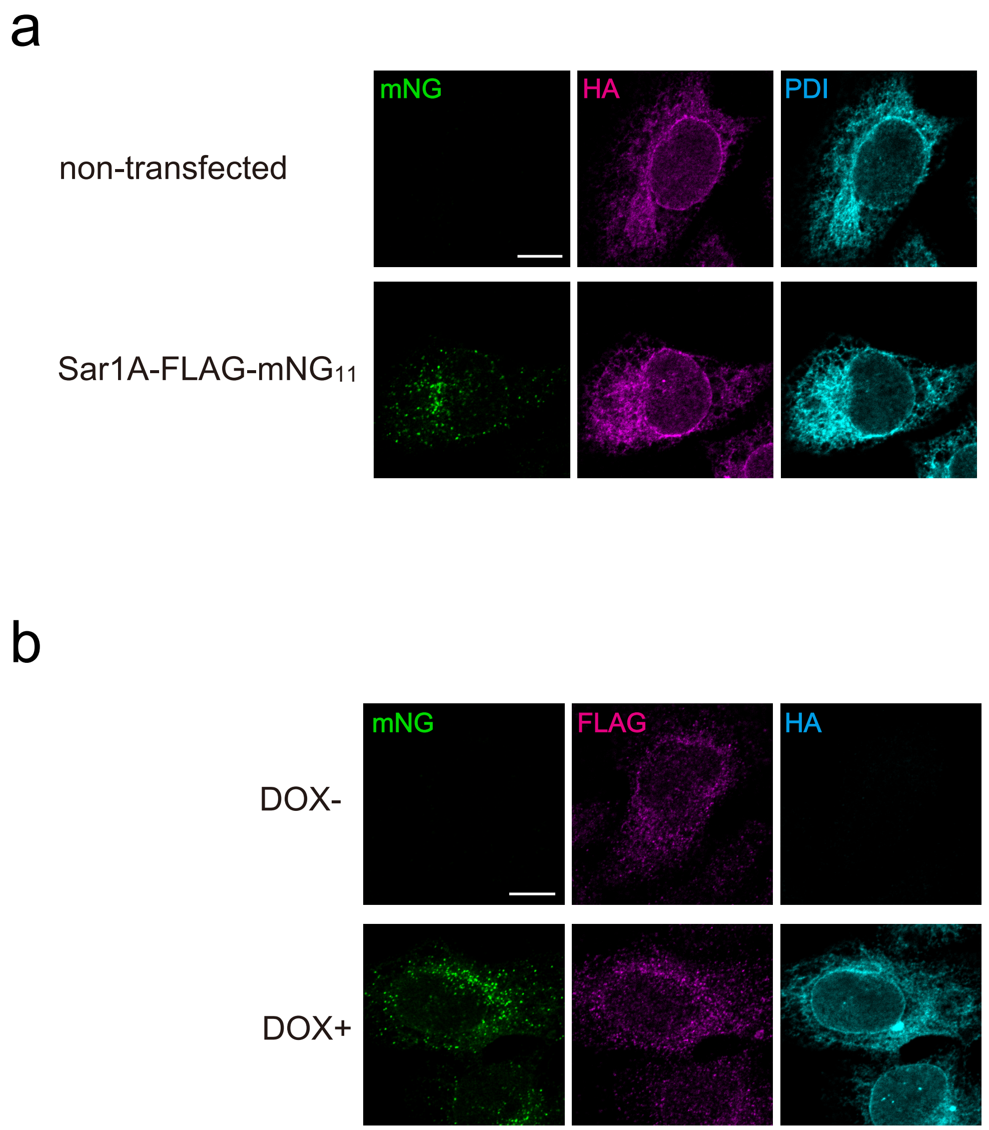


**Supplementary Fig. 1: Validation of SAIYAN technology.**

**a** Doxycycline-inducible HeLa cells expressing the membrane-spanning region of TANGO1S and HA-tag fused to 10 of the 11 bundles of mNG (HA-mNG1-10 cells) were either non-transfected or transfected with Sar1A constructs with a FLAG tag and a glycine linker fused to the 11th bundle of mNG (Sar1A-FLAG-mNG11). The cells were fixed and stained with anti-HA and anti-PDI antibodies. Scale bars = 10 μm. **b** HA-mNG1-10 cells, treated with or without doxycycline, were transfected with Sar1A-FLAG-mNG11. The cells were fixed and stained with anti-HA and anti-FLAG antibodies. Scale bars = 10 μm.

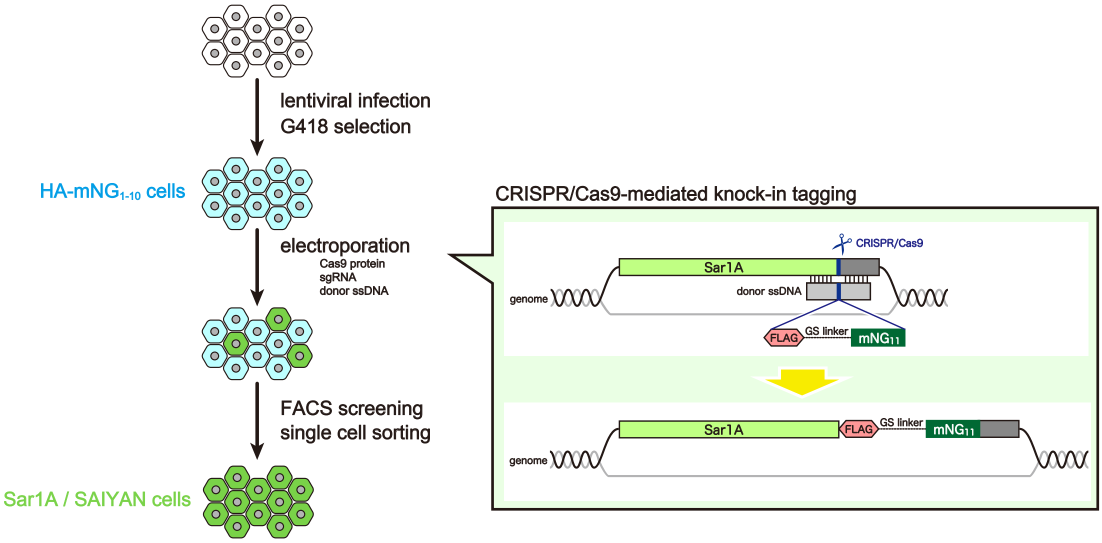


**Supplementary Fig. 2: Schematic diagram illustrating the process of construction of Sar1A/SAIYAN cells.**

Doxycycline-inducible stable cell lines expressing TM-mNG1-10 were established using a lentiviral system and G418 selection (HA-mNG1-10 cells). Stable cells were subsequently electroporated with Cas9 protein, sgRNA, and ssDNA to facilitate the knock in of FLAG-mNG11 into the Sar1A locus of the genome. Cells were treated with doxycycline for 24 h and further sorted by FACS to isolate single cells exhibiting mNG signals into 96 well plates. The expanded cell population was then collected and subjected to genomic sequencing. Positive clones were identified and selected for further analysis (Sar1A/SAIYAN cells).

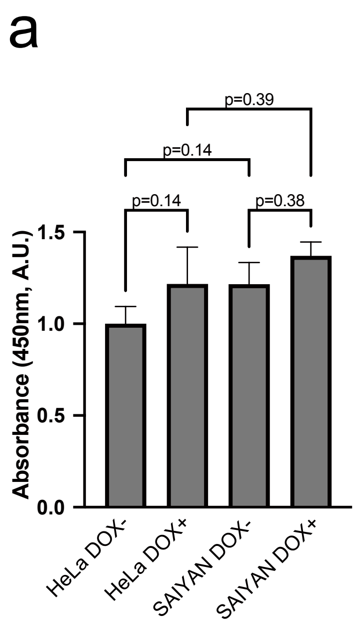


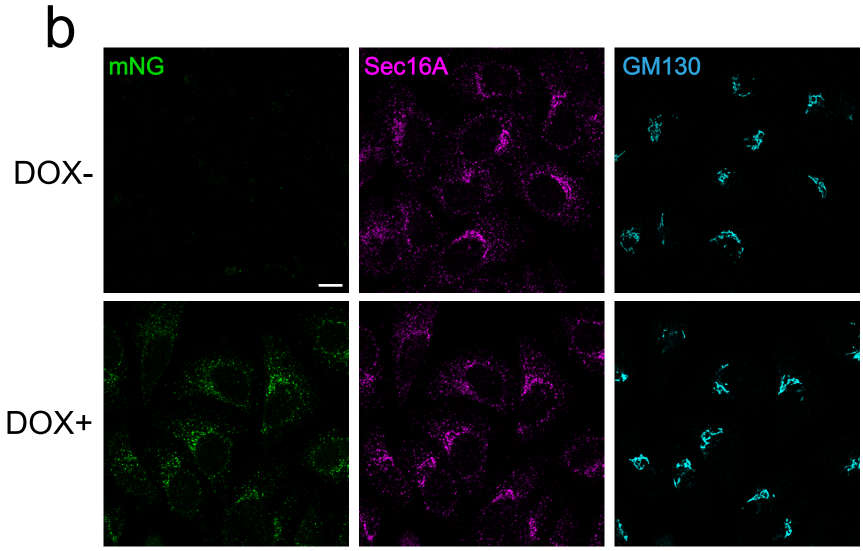


**Supplementary Fig. 3: Sar1A/SAIYAN (HeLa) cells proliferate normally and exhibit a regular structure of the Golgi apparatus.**

**a** HeLa and Sar1A/SAIYAN (HeLa) cells were treated with or without doxycycline for 24 h and cell viability was measured and normalized using untreated HeLa cells as control. Error bars represent the means ± SEM. **b** Sar1A/SAIYAN (HeLa) cells, treated with or without doxycycline, were fixed and stained with anti-Sec16-C and anti-GM130 antibodies. Scale bars = 10 μm.

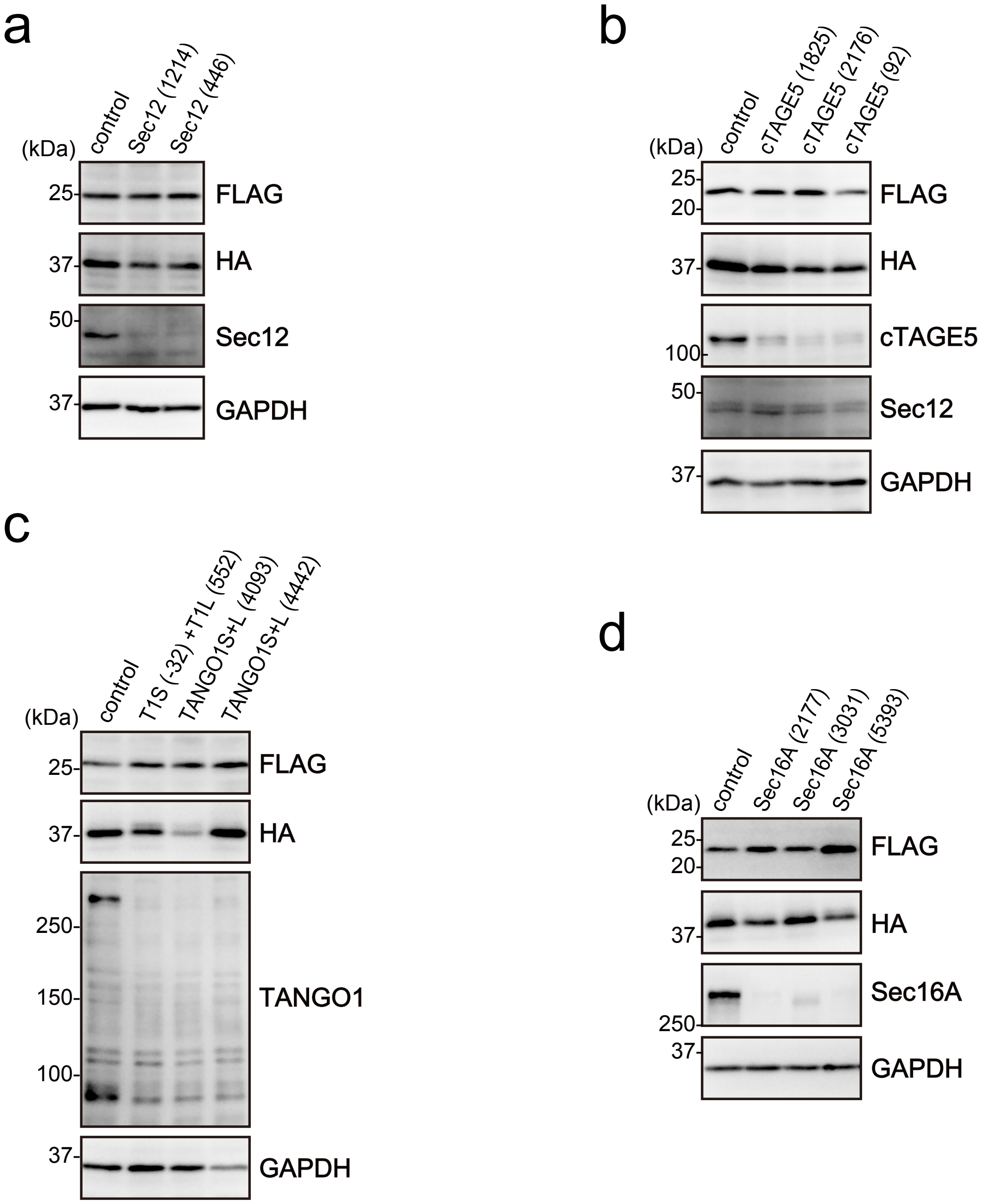


**Supplementary Fig. 4: Validation of knockdown efficiency using western blotting for Fig. 2.**

**a** Sar1A/SAIYAN (HeLa) cells transfected with the indicated siRNAs were lysed, and subjected to SDS-PAGE, followed by western blotting with anti-FLAG, anti-HA, anti-Sec12, and anti-GAPDH antibodies. **b** Sar1A/SAIYAN (HeLa) cells transfected with the indicated siRNAs were lysed, and subjected to SDS-PAGE, followed by western blotting with anti-FLAG, anti-HA, anti-cTAGE5 CC1, anti-Sec12 and anti-GAPDH antibodies. **c** Sar1A/SAIYAN (HeLa) cells transfected with the indicated siRNAs were lysed, and subjected to SDS-PAGE, followed by western blotting with anti-FLAG, anti-HA, anti-TANGO1 CC1, and anti-GAPDH antibodies. **d** Sar1A/SAIYAN (HeLa) cells transfected with the indicated siRNAs were lysed, and subjected to SDS-PAGE followed by western blotting with anti-FLAG, anti-HA, anti-Sec16-N, and anti-GAPDH antibodies.

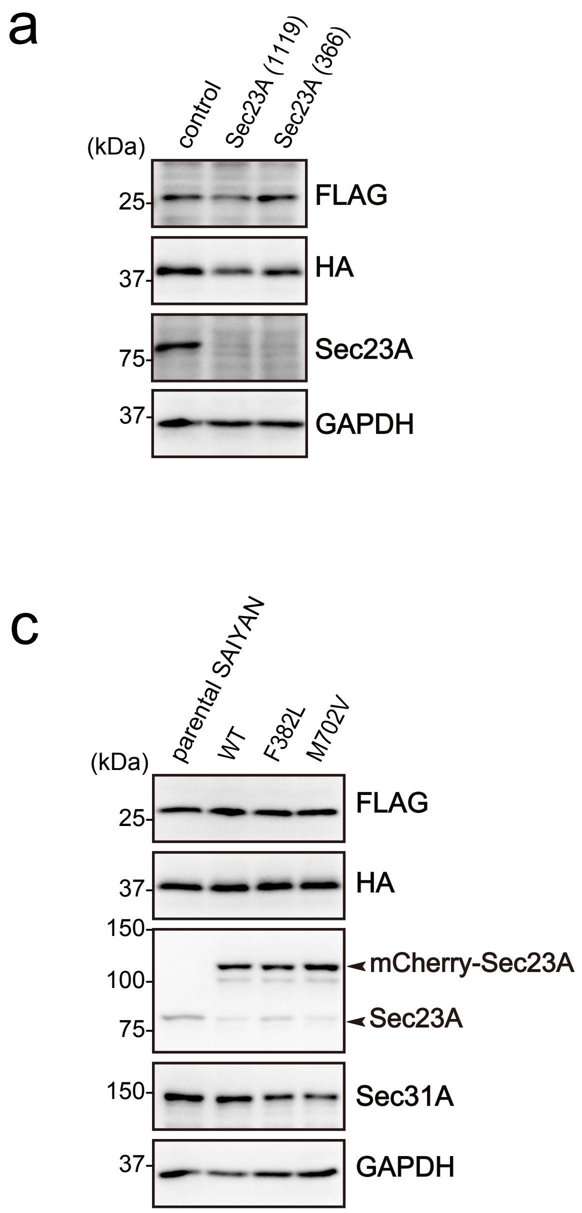


**Supplementary Fig. 5: Validation of knockdown efficiency using western blotting for Fig. 3.**

**a** Sar1A/SAIYAN (HeLa) cells transfected with the indicated siRNAs were lysed, and subjected to SDS-PAGE, followed by western blotting with anti-FLAG, anti-HA, anti-Sec23A, and anti-GAPDH antibodies. **b** Sar1A/SAIYAN (HeLa) cells transfected with the indicated siRNAs were lysed, and subjected to SDS-PAGE, followed by western blotting with anti-FLAG, anti-HA, anti-Sec31A (rabbit) and anti-GAPDH antibodies. **c** Sar1A/SAIYAN (HeLa) cells were stably expressed using mCherry-tagged Sec23A constructs as indicated. Cells were lysed, and subjected to SDS-PAGE, followed by western blotting with anti-FLAG, anti-HA, anti-Sec23A, anti-Sec31A (rabbit), and anti-GAPDH antibodies.

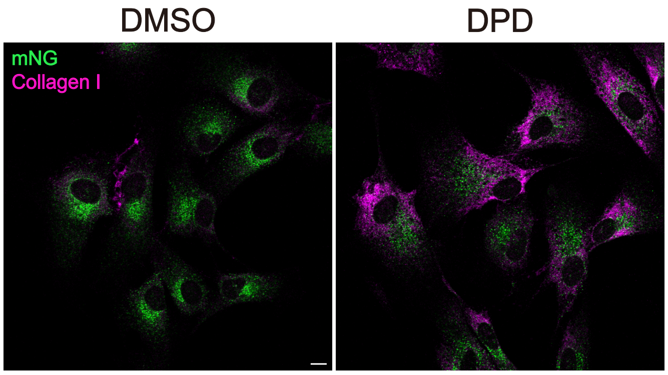


**Supplementary Fig. 6: DPD treatment accumulates collagen I within the ER of Sar1A/SAIYAN (BJ-5ta) cells.**

Sar1A/SAIYAN (BJ-5ta) cells were treated with DMSO or 0.5 mM DPD and incubated for 16 h. Cells were fixed and stained with an anti-collagen I antibody. Scale bars = 10 μm.

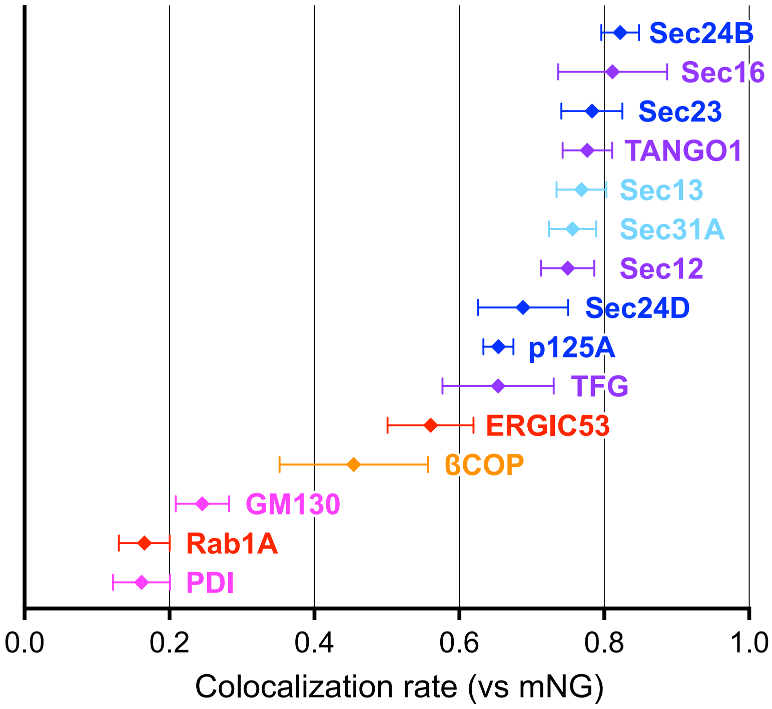


**Supplementary Fig. 7: Quantification of Pearson's correlation coefficient to quantify the degree of colocalization in Sar1A/SAIYAN (HeLa) cells.**

Sar1A/SAIYAN (HeLa) cells were fixed and stained with anti-Sec16-C, anti-ERGIC53, anti-Sec23, anti-Sec24B, anti-Sec24D, anti-p125A, anti-TANGO1-CT, anti-Sec12, anti-TFG, anti-Sec13, anti-Sec31A (mouse), anti-β-COP, anti-GM130, anti-PDI, and anti-Rab1A antibodies. Images were captured using the Airyscan2. n=5. cyan; outer COPII coats, blue; inner COPII coats, purple; endoplasmic reticulum (ER) exit site resident proteins, red; ERGIC proteins, orange; COPI protein, magenta; ER and Golgi proteins. Error bars represent the mean 95% CI.

**Supplementary Table 1: siRNA sequences used in this study**

| oligo | sense | antisense |
|---|---|---|
| cTAGE5 (1825) | CCG CCA GGA CAA UCA UAU CCU GAU U | AUC AGG AUA UGA UUG UCC UGG CGG |
| cTAGE5(2176) | GCC AUG UUU GGA GCU UCU CGA GAU U | AAU CUC GAG AAG CUC CAA ACA UGG C |
| cTAGE5 (92) | GAC CAG AUU CUA AUC UUU AUG GUU U | AAA CCA UAA AGA UUA GAA UCU GGU C |
| Sar1A (269) | UCC CAG CAA UUA AUG GGA UUG UCU U | AAG ACA AUC CCA UUA AUU GCU GGG A |
| Sar1A (237) | CGA GCA AGC ACG UCG CGU UUG GAA A | UUU CCA AAC GCG ACG UGC UUG CUC G |
| Sec12 (1214) | CCA UCC UGC UGC UCC AGA GUG CCU U | AAG GCA CUC UGG AGC AGC AGG AUG G |
| Sec12 (446) | CAG ACU UUA GCU CCG AUC CAC UGC A | UGC AGU GGA UCG GAG CUA AAG UCU G |
| Sec16A (2177) | GGG CGC AAA GUG AGC UGC CAG AUU U | AAA UCU GGC AGC UCA CUU UGC GCC C |
| Sec16A (3031) | CCG UCC CAU UCU GAC AGC CUC GCU U | AAG CGA GGC UGU CAGAAU GGG ACG G |
| Sec16A (5393) | CCC UGC CUA GUU UCC AGG UGU UUA A | UUA AAC ACC UGG AAA CUA GGC AGG G |
| Sec23A (1119) | GGG UGA UUC UUU CAA UAC UUC CUU A | UAA GGA AGU AUU GAA AGA AUC ACC C |
| Sec23A (366) | GCG UGG UCC UCA GAU GCC UUU GAU A | UAU CAA AGG CAU CUG AGG ACC ACG C |
| Sec31A (1805) | CCA UAG CAG GUG GAC AAG AAC UCU U | AAG AGU UCU UGU CCA CCU GCU AUG G |
| Sec31A (3297) | CCA GGC CAA UAA GCU GGG UGU CUA A | UUA GAC ACC CAG CUU AUU GGC CUG G |
| TANS (-32) | GAA UUG UCG CUU GCG UUC AGC UGU U | AAC AGC UGA ACG CAA GCG ACA AUU C |
| TANL (552) | CAA CUC AGA GGA AAG UGA UAG UGU A | UAC ACU AUC ACU UUC CUC UGA GUU G |
| TANS+L (4093) | CAG GAA AUC GAA GAC UGG AGU AAA U | AUU UAC UCC AGU CUU CGA UUU CCU G |
| TANS+L (4442) | CCG UGU CCA CUA AAU GUA ACC UGG A | UCC AGG UUA CAU UUA GUG GAC ACG G |
